# An Adaptive Optimizer without User-Specified Hyperparameters through Historical Projection and Self-Calibrated Scaling

**DOI:** 10.1101/348557

**Authors:** Kazunori D Yamada

## Abstract

Adaptive optimizers reduce sensitivity to coordinate-wise gradient scale, but they usually still require the user to choose an overall learning rate and often a schedule. We introduce YamAdam, an adaptive optimizer that requires no user-specified optimizer hyperparameters. YamAdam combines short-memory Adam moments with both decay factors fixed at 1*/*2, a projection that removes only the historical component opposing the current mini-batch gradient, a one-time calibration of a reference update radius, and subsequent scaling based on dimensionless directional statistics. We evaluate YamAdam without user-specified optimizer settings on six workloads spanning image classification, image reconstruction, tabular classification and regression, and language modeling. The comparison contains 30 fixed settings from 10 optimizer families, with every setting reported separately. Our primary measure, Δ_TTT_, is the comparator-minus-YamAdam difference in budget-normalized time to five predefined targets, averaged equally across workloads. All 30 comparator settings produced a positive Δ_TTT_, ranging from +0.003526 to +0.210524. Across 540 setting– workload–seed comparisons, YamAdam was faster in 504, tied in 14, and slower in 22. Across 2,700 target observations, both methods reached 2,634 targets and only YamAdam reached 66; no target was reached only by the comparator or by neither method. These results show that an optimizer requiring no user-specified optimizer hyperparameters can attain predefined targets rapidly across heterogeneous workloads.

## 1 Introduction

Adam and AdamW are widely used because they adapt each coordinate to recent gradient scale [1, 2]. They nevertheless leave an overall learning rate, and often a schedule, to the user. Selecting those values separately for every workload creates a second optimization problem outside model training and makes conclusions depend on the tuning budget.

Several methods reduce this burden. D-Adaptation and Prodigy estimate a global scale, DoG uses the distance traveled from initialization, Mechanic rescales a base optimizer, and Schedule-Free methods remove an externally specified schedule [3–7]. These methods demonstrate that substantial parts of learning-rate selection can be automated. They also use different internal rules and expose different controls. Moreover, fully parameter-free stochastic optimization is impossible under some oracle models [8].

We therefore ask a practical question: can the same adaptive optimizer reach useful targets rapidly across heterogeneous workloads without workload-specific optimizer hyperparameter tuning?

We propose Yamada modified Adam (YamAdam). It makes two decisions at every update: which direction to move and how far to move. The direction starts from a short-memory Adam estimate. If the part inherited from earlier gradients opposes the current gradient, only that opposing part is removed; the contribution of the current gradient is retained. The first nonzero update uses the current mini-batch to calibrate a reference distance. Later distances are computed from dimensionless directional statistics and require no further trial-point loss evaluations.

The central contribution of this work is an adaptive optimizer that combines a projection acting only on the historical component, scale calibration at the first update, and subsequent scale adjustment independent of loss units into a single update rule, thereby determining both direction and update distance without user-specified optimizer hyperparameters.

## 2 Related work

Adam combines first and second moments, bias correction, and coordinate-wise normalization [1]. NAdam adds Nesterov-style momentum, and AdamW decouples weight decay from the adaptive update [2, 9]. SMORMS3 uses a rapidly adapting per-parameter memory [10]. YamAdam uses Adam moments with both decay factors fixed at 1*/*2; its main changes concern the direction formed from those moments and the global distance moved along it.

Methods that reduce learning-rate or schedule specification include D-Adaptation, Prodigy, DoG, Mechanic, and Schedule-Free optimization [3–7]. In contrast, YamAdam calibrates its scale at the first update and subsequently adjusts its update distance from internal quantities that are independent of loss units.

The relation between a current gradient and accumulated motion has been used for restart, update rejection, masking, sign replacement, and momentum-gradient policies [11–15]. GEM and PCGrad project gradients to handle conflicts across tasks [16, 17]. YamAdam instead uses one current mini-batch gradient and corrects only the historical part of a preconditioned proposal.

Armijo backtracking and stochastic line search select a step length by evaluating trial points [18, 19]. L4 also uses loss information to adapt the step repeatedly [20]. YamAdam uses trial losses only during its first nonzero update; subsequent distances follow a deterministic recurrence.

## 3 Method

### 3.1 The two decisions made at each update

Every optimizer update must answer two questions: in which direction should the parameters move, and how far should they move in that direction? YamAdam answers them separately. It obtains a direction from recent gradients, removes only the part of the stored history that conflicts with the current gradient, and then gives that direction unit length. It determines the overall distance once from the loss on the first mini-batch and adjusts later distances by comparing the current directional signal with its own past values.

We use *k* = 1, 2, … for completed nonzero optimizer updates. Immediately before update *k, θ*_*k*− 1_ is the complete parameter vector, *ℓ*_*k*_(*θ*) is the loss of the current mini-batch evaluated at parameters *θ*, and

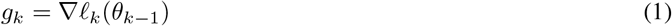

is its gradient at the current parameters. Inner products such as ⟨*x, y*⟩ mean the sum of coordinate-wise products, and ∥*x*∥_2_ is the Euclidean length of a vector. Only coordinates for which the model computes a derivative participate in an update. A parameter reported by automatic differentiation as having no gradient is left unchanged and is omitted from that update’s inner products and norms.

The letter *d*_*k*_ denotes the direction proposal before it is given unit length, *w*_*k*_ = *d*_*k*_*/* ∥*d*_*k*_∥_2_ its unit-length form, *C* the reference distance determined on the first update, and *ρ*_*k*_ the distance actually used on update *k*. A bar denotes an uncorrected moving average, a tilde a vector corrected by projection, and a hat a vector rescaled to unit length. Vector notation keeps the equations readable; the implementation keeps each parameter in its original tensor shape.

### 3.2 Short-memory Adam proposal

Let *t* be the number of consecutive updates for which a particular parameter has supplied a gradient, and let *g*_*t*_ denote that parameter’s gradient at its *t*th consecutive participating update. Starting from 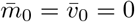, YamAdam gives half of the moving-average weight to the current gradient and half to the previous average:

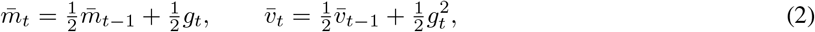

The usual bias correction removes the initial bias toward zero:

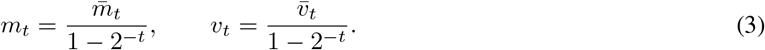

These are exactly Adam’s bias-corrected first and second moments with both decay factors fixed at 1*/*2. The short memory makes old gradients lose influence quickly. We also define *m*_0_ = *v*_0_ = 0.

The corrected first moment can be written as a contribution from the previous corrected moment plus a contribution from the current gradient. Their weights are

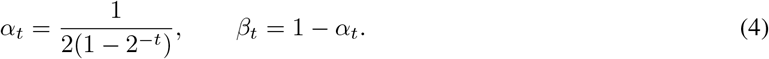

The factor *α*_*t*_ is not an additional design choice. It follows from expanding the bias-corrected recurrence in Eqs. (2)–(3). It equals 1 at the first observation, then 2*/*3, 4*/*7, and approaches 1*/*2; hence the current gradient supplies all of the first direction and about half of later directions.

For each coordinate, define the common scale

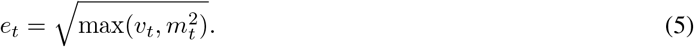

The maximum is taken coordinate by coordinate. Dividing both contributions by this same current scale gives

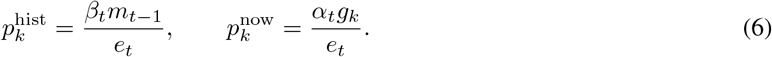

Here 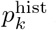 is the part inherited from earlier gradients and 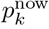 is the part supplied by the current gradient. Because they use the same denominator, their sum is the ordinary normalized Adam first moment with decay factors 1*/*2. If *e*_*t*_ = 0, both parts are defined as zero. If a parameter leaves the active set while other parameters complete an update, its local count and moments restart when it becomes active again.

### 3.3 Correcting only opposing history

Momentum can keep pointing along an old direction even after the current gradient changes. Projecting the complete Adam proposal would also alter the new information just supplied by the current gradient. YamAdam therefore modifies only 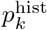:

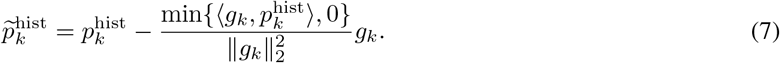

The sign of 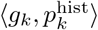 tells whether the stored part helps the present descent direction. A nonnegative value leaves the history unchanged. A negative value removes exactly its component parallel to −*g*_*k*_ and keeps the perpendicular component. The vectors whose inner product with *g*_*k*_ is nonnegative form one side of a flat boundary through the origin, called a half-space. Equation (7) is the smallest Euclidean change that places the historical vector in that half-space.

The current-gradient contribution is then added without modification:

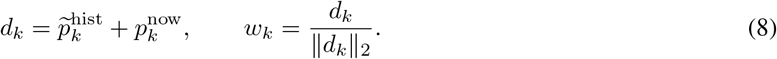

The parameter step will be −*w*_*k*_. Because the retained current contribution has positive alignment with *g*_*k*_, −*w*_*k*_ is a local descent direction for the current mini-batch whenever the gradient is nonzero.

### 3.4 Calibrating the initial distance

A unit vector determines a direction, but it does not say how far to move in parameter space. YamAdam adapts the Armijo sufficient-decrease condition [18] into a one-time automatic calibration. Ordinary Armijo search chooses a step repeatedly. Here trial losses are used only on the first nonzero update; the accepted distance becomes a reference for all later updates.

The first gradient is initially converted into an auxiliary probe. Let *µ* be the mean squared coordinate of *g*_1_, let *q* be the coordinate-bounded version of that gradient, and let *q* be *q* rescaled to unit length:

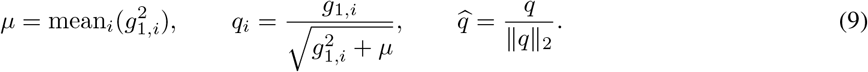

The probe is used only to find a plausible distance. Bounding each coordinate prevents one exceptionally large gradient coordinate from controlling that search; it does not replace the actual update direction *w*_1_.

For a unit direction *u*, a trial distance *R* is accepted when

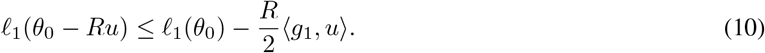

The right-hand side is the decrease predicted by the current gradient, with a fixed factor 1*/*2 allowing for local curvature. The same mini-batch and stochastic state are replayed at every trial. Starting from *R* = 1, the search halves a rejected distance until it is accepted. If the first distance is accepted and a finite upper boundary can be sought, it doubles the distance until the next finite trial is rejected and retains the preceding accepted distance. If parameter rounding makes a trial produce no change, the distance is enlarged without evaluating the loss. If no finite upper boundary exists along a locally linear or concave ray, the rule returns to the first accepted distance instead of using an arbitrarily large value.

The distance found with *q* is then checked along the actual direction *w*_1_ and halved if needed. The final accepted value, *C*, is both the length of the first update and the fixed reference for later updates. Thus the calibration uses the Armijo idea, but differs from ordinary repeated Armijo search in two ways: it first searches along a bounded auxiliary direction and rechecks the result along the actual direction, and it is performed only once.

### 3.5 Scaling later updates

Later updates do not evaluate trial losses. Instead, the current gradient predicts the local loss decrease per unit movement along −*w*_*k*_. This first-order prediction is

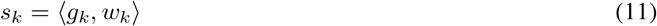

because *ℓ*_*k*_(*θ*_*k*− 1_ − *ρw*_*k*_) ≈ *ℓ*_*k*_(*θ*_*k*− 1_) − *ρs*_*k*_ for a small distance *ρ*. Thus “current descent strength” means *s*_*k*_: it is a local prediction, not an observed loss decrease after the update. To compare it with earlier predictions, YamAdam divides it by their running root-mean-square:

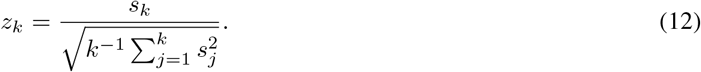

The dimensionless value *z*_*k*_ says whether the present predicted decrease per unit distance is strong or weak relative to its own history. Multiplying all mini-batch losses by the same positive constant multiplies both the numerator and denominator and therefore leaves *z*_*k*_ unchanged.

Two clocks then propose distances:

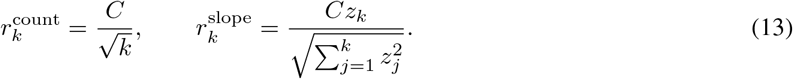

The count clock 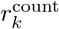 decreases as 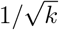. The slope clock 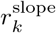 becomes relatively larger when the present directional prediction is strong compared with its history and smaller when it is weak. Both are expressed in units of the initially calibrated distance *C*.

To prevent either clock from overwhelming the other, define the factor-two combiner

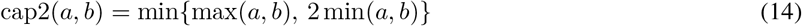

It selects the larger positive value unless that value exceeds twice the smaller one, in which case it returns twice the smaller one. The baseline distance is

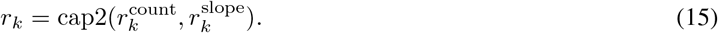

The two clocks normally decrease even if a previously conflicting historical direction later becomes useful again. YamAdam therefore permits a limited recovery of distance, but only after evidence of both events: a projection has occurred at some earlier update, and the historical component agrees with the current gradient on two consecutive updates. The first condition shows that stale history was once harmful; the second shows that the retained history is now repeatedly useful. Current agreement is measured by

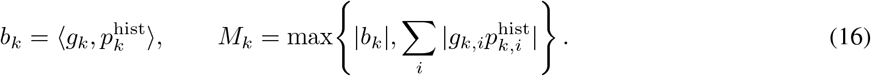

where *b*_*k*_ is the signed alignment and *M*_*k*_ is a cancellation-safe scale for that inner product. If no projection is needed and *M*_*k*_ *>* 0, let *a*_*k*_ = clip(*b*_*k*_*/M*_*k*_, 0, 1); otherwise let *a*_*k*_ = 0. The function clip(*x*, 0, 1) limits a value to the interval from zero to one. Hence *a*_*k*_ is a dimensionless agreement score, not a gradient magnitude.

Let *h*_*k*− 1_ = 1 if any projection occurred before update *k* and zero otherwise, and set *a*_0_ = 0. The gate *ξ*_*k*_ = *h*_*k* − 1_**1** {*a*_*k* −1_ *>* 0} is one only when an earlier conflict and positive agreement on the preceding update have both been observed. The indicator **1**{·} equals one when its condition is true and zero otherwise. The largest permitted recovered distance is

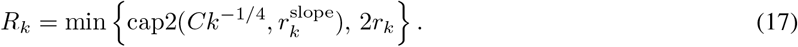

It cannot exceed twice the baseline *r*_*k*_ and still decreases no more slowly than *Ck*^−1*/*4^. With [*x*]_+_ = max(*x*, 0), the recovered distance is

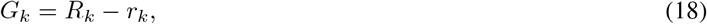

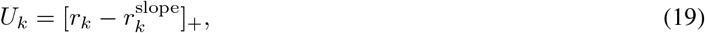

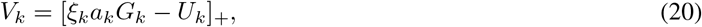

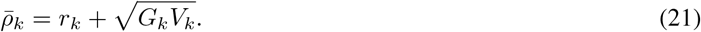

Here *G*_*k*_ is the unused room below the ceiling. The term *U*_*k*_ charges the amount by which the baseline already exceeds the slope clock. The remaining support *V*_*k*_ is positive only when the earlier-conflict gate, the preceding agreement, and the current agreement jointly outweigh that charge. The geometric mean 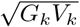 then releases only part of the available room. Consequently, no recovery occurs without the stated evidence and 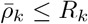.

One additional lower bound is available only when the initial calibration backtracked. In that case, the accepted distance *C* has an adjacent rejected dyadic trial at 2*C*. This observed boundary is converted to a coefficient using the first unnormalized proposal:

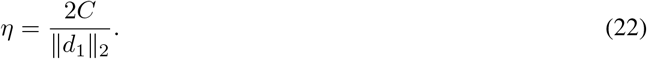

The applied distance is then

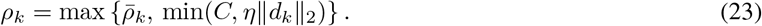

The floor min(*C*, η∥*d*_*k*_∥_2_) prevents the distance from becoming very small while the raw direction proposal remains large, and the cap at *C* keeps it within the initial reference. When *C* was determined on the auxiliary probe, applying the coefficient to *d*_1_ is an extrapolation; when it was determined by the final check, the rejected point lay on the actual first direction. Neither case guarantees later loss decrease. If the calibration did not backtrack, no such coefficient is created and 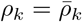. The final update is

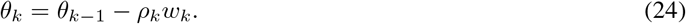

Ignoring rounding error, the update rule has three useful properties. First, for a nonzero gradient, ⟨*g*_*k*_, *w*_*k*_⟩ *>* 0, so the actual step direction −*w*_*k*_ is a descent direction for the current mini-batch. This follows because an opposing historical component is removed and the current-gradient contribution is retained. Second, every applied distance is positive and no greater than the calibrated reference *C*. Third, the parameter trajectory is invariant under a positive affine transformation of the loss, 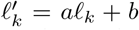 with fixed *a >* 0 and fixed *b*. This means that changing the unit or zero point of the same loss does not change the updates; it does not make different loss functions equivalent. These properties describe the update rule rather than prove convergence, and later updates are not checked by line search.

### 3.6 Numerical implementation

The equations above state the exact-arithmetic rule. Three fixed safeguards make that rule reproducible on finite-precision hardware. First, the maximum in Eq. (5) is coordinate-wise. Exact weighted moments satisfy 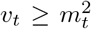, so it ordinarily returns *v*_*t*_; retaining 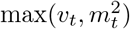 prevents a variance-like discrepancy caused by rounding. Second, projection uses the positive margin

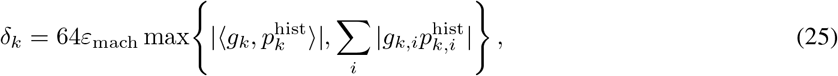

where *ε*_mach_ is the machine epsilon of the gradient’s floating-point type. The projected history is placed on the positive side by this margin. For the agreement score, the implementation sets *a*_*k*_ = 0 when *b*_*k*_ ≤ *δ*_*k*_ and otherwise clips (*b*_*k*_ − *δ*_*k*_)*/M*_*k*_ to [0, 1]. Thus numerical noise alone does not count as positive agreement. The factor 64 is a fixed rounding-error allowance, not a user setting. Third, each part of the initial search is limited to 64 trials to prevent an endless search; this separate 64 is also fixed. Trial evaluations replay the same mini-batch, random-number state, and mutable model buffers. These safeguards and their exception handling are implemented in the public code.

## 4 Experimental protocol

### 4.1 Workloads and paired runs

We evaluate six workloads spanning image classification, image reconstruction, tabular classification and regression, and language modeling (Table 1). Dataset names identify data sources rather than canonical training protocols. No test split is used. Validation is recorded before training and after every optimizer update. Within each paired workload–seed comparison, initialization, data order, augmentation, numerical precision, and update budget are identical.

**Table 1:**
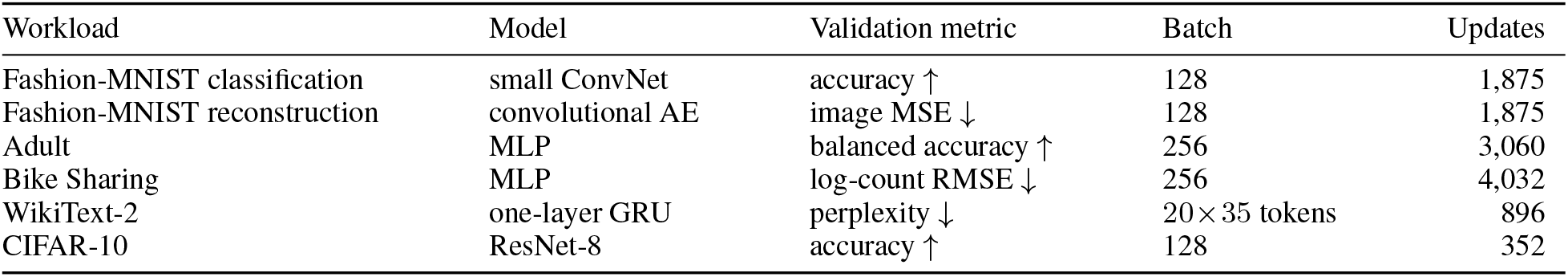
Workloads and fixed update budgets. These bounded budgets are not claims of training to convergence.

We use Fashion-MNIST [21], the Adult and Bike Sharing records from the UCI Machine Learning Repository [22, 23], WikiText-2 [24], and the CIFAR-10 training batches [25]. The language model uses a one-layer GRU [26]; the image classifier uses a small residual network with group normalization [27, 28]. Experiments use PyTorch, FP32 arithmetic, deterministic algorithms, and seeds 30–32. The primary speed axis is the number of completed optimizer updates, not wall-clock time.

### 4.2 Fixed comparator settings

The comparison contains three fixed settings for each of 10 optimizer families: D-Adaptation Adam, AdamW, NAdamW, Prodigy, Schedule-Free AdamW, Mechanic with Adam, Cautious AdamW, Grams, SMORMS3, and DoG. Every setting is reported as its own row. We do not create a tuned row, family winner, oracle, workload-specific selection, or best-of-grid result. AdamW and NAdamW use learning rates {10^−4^, 10^−3^, 10^−2^}. Table 2 shows the field varied across the three rows of each family. Other fixed values, the implementation and version used, data splits, preprocessing, and model definitions are provided in the public repository.

**Table 2:**
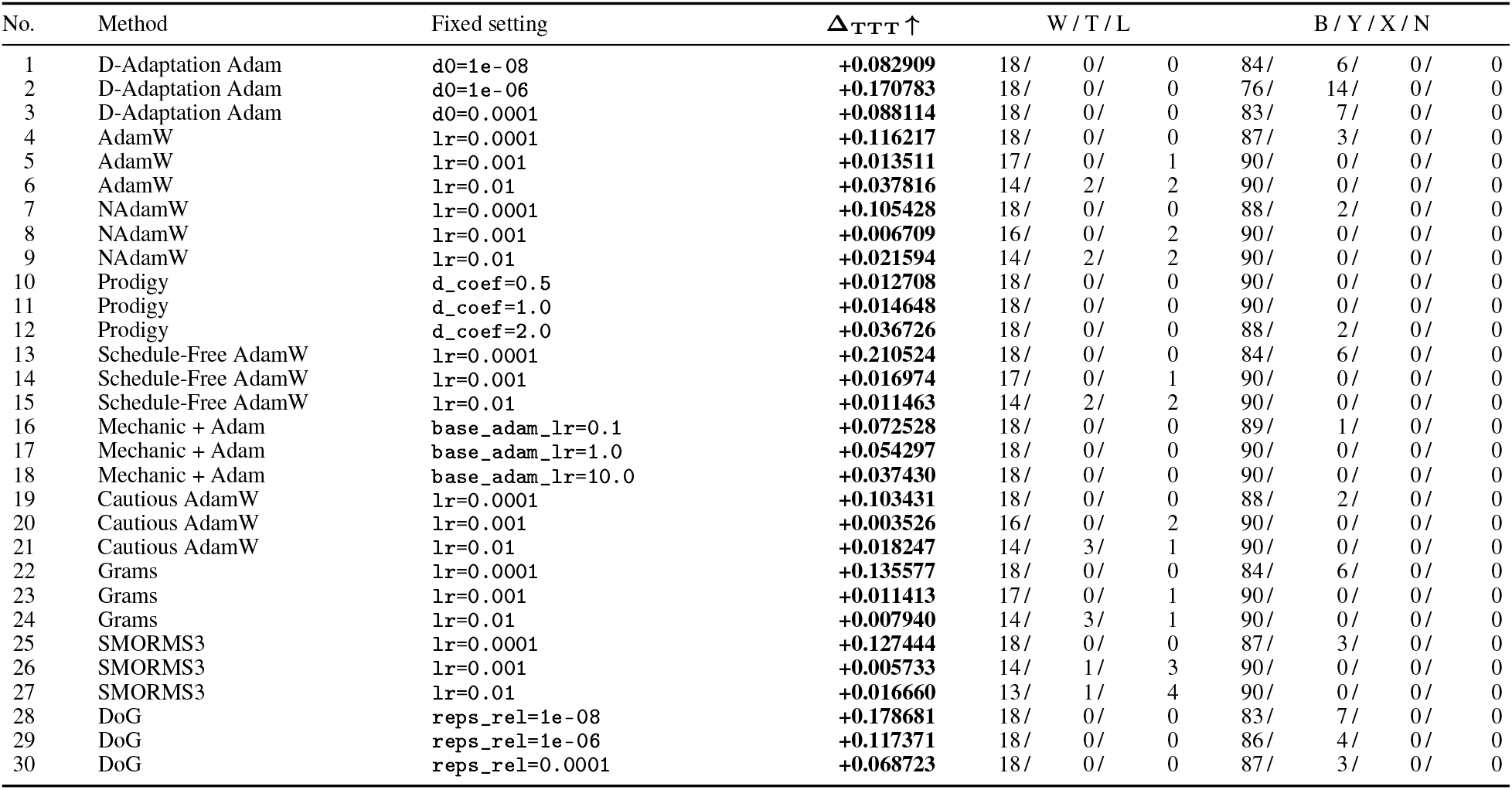
Primary comparison with 30 fixed settings. Δ_TTT_ is the comparator-minus-YamAdam difference in budget-normalized restricted target time; positive values favor YamAdam. W / T / L counts YamAdam wins, ties, and losses across 18 workload–seed cases. B / Y / X / N counts both reached, YamAdam only, comparator only, and neither reached across 90 targets.

| No. | Method | Fixed setting | $\Delta_{\text{TTT}} \uparrow$ | W / T / L | | | B / Y / X / N | | | |
| --- | --- | --- | --- | --- | --- | --- | --- | --- | --- | --- |
| 1 | D-Adaptation Adam | d0=1e-08 | **+0.082909** | 18/ | 0/ | 0 | 84/ | 6/ | 0/ | 0 |
| 2 | D-Adaptation Adam | d0=1e-06 | **+0.170783** | 18/ | 0/ | 0 | 76/ | 14/ | 0/ | 0 |
| 3 | D-Adaptation Adam | d0=0.0001 | **+0.088114** | 18/ | 0/ | 0 | 83/ | 7/ | 0/ | 0 |
| 4 | AdamW | lr=0.0001 | **+0.116217** | 18/ | 0/ | 0 | 87/ | 3/ | 0/ | 0 |
| 5 | AdamW | lr=0.001 | **+0.013511** | 17/ | 0/ | 1 | 90/ | 0/ | 0/ | 0 |
| 6 | AdamW | lr=0.01 | **+0.037816** | 14/ | 2/ | 2 | 90/ | 0/ | 0/ | 0 |
| 7 | NAdamW | lr=0.0001 | **+0.105428** | 18/ | 0/ | 0 | 88/ | 2/ | 0/ | 0 |
| 8 | NAdamW | lr=0.001 | **+0.006709** | 16/ | 0/ | 2 | 90/ | 0/ | 0/ | 0 |
| 9 | NAdamW | lr=0.01 | **+0.021594** | 14/ | 2/ | 2 | 90/ | 0/ | 0/ | 0 |
| 10 | Prodigy | d_coef=0.5 | **+0.012708** | 18/ | 0/ | 0 | 90/ | 0/ | 0/ | 0 |
| 11 | Prodigy | d_coef=1.0 | **+0.014648** | 18/ | 0/ | 0 | 90/ | 0/ | 0/ | 0 |
| 12 | Prodigy | d_coef=2.0 | **+0.036726** | 18/ | 0/ | 0 | 88/ | 2/ | 0/ | 0 |
| 13 | Schedule-Free AdamW | lr=0.0001 | **+0.210524** | 18/ | 0/ | 0 | 84/ | 6/ | 0/ | 0 |
| 14 | Schedule-Free AdamW | lr=0.001 | **+0.016974** | 17/ | 0/ | 1 | 90/ | 0/ | 0/ | 0 |
| 15 | Schedule-Free AdamW | lr=0.01 | **+0.011463** | 14/ | 2/ | 2 | 90/ | 0/ | 0/ | 0 |
| 16 | Mechanic + Adam | base_adam_lr=0.1 | **+0.072528** | 18/ | 0/ | 0 | 89/ | 1/ | 0/ | 0 |
| 17 | Mechanic + Adam | base_adam_lr=1.0 | **+0.054297** | 18/ | 0/ | 0 | 90/ | 0/ | 0/ | 0 |
| 18 | Mechanic + Adam | base_adam_lr=10.0 | **+0.037430** | 18/ | 0/ | 0 | 90/ | 0/ | 0/ | 0 |
| 19 | Cautious AdamW | lr=0.0001 | **+0.103431** | 18/ | 0/ | 0 | 88/ | 2/ | 0/ | 0 |
| 20 | Cautious AdamW | lr=0.001 | **+0.003526** | 16/ | 0/ | 2 | 90/ | 0/ | 0/ | 0 |
| 21 | Cautious AdamW | lr=0.01 | **+0.018247** | 14/ | 3/ | 1 | 90/ | 0/ | 0/ | 0 |
| 22 | Grams | lr=0.0001 | **+0.135577** | 18/ | 0/ | 0 | 84/ | 6/ | 0/ | 0 |
| 23 | Grams | lr=0.001 | **+0.011413** | 17/ | 0/ | 1 | 90/ | 0/ | 0/ | 0 |
| 24 | Grams | lr=0.01 | **+0.007940** | 14/ | 3/ | 1 | 90/ | 0/ | 0/ | 0 |
| 25 | SMORMS3 | lr=0.0001 | **+0.127444** | 18/ | 0/ | 0 | 87/ | 3/ | 0/ | 0 |
| 26 | SMORMS3 | lr=0.001 | **+0.005733** | 14/ | 1/ | 3 | 90/ | 0/ | 0/ | 0 |
| 27 | SMORMS3 | lr=0.01 | **+0.016660** | 13/ | 1/ | 4 | 90/ | 0/ | 0/ | 0 |
| 28 | DoG | reps_rel=1e-08 | **+0.178681** | 18/ | 0/ | 0 | 83/ | 7/ | 0/ | 0 |
| 29 | DoG | reps_rel=1e-06 | **+0.117371** | 18/ | 0/ | 0 | 86/ | 4/ | 0/ | 0 |
| 30 | DoG | reps_rel=0.0001 | **+0.068723** | 18/ | 0/ | 0 | 87/ | 3/ | 0/ | 0 |

YamAdam is run without user-specified optimizer settings on six workloads and three seeds, giving 18 YamAdam trajectories. The 30 comparator settings give 540 trajectories. The public comparison therefore contains 558 unique training runs, 540 paired setting–workload–seed cases, and 2,700 setting–workload–seed–target observations.

### 4.3 Targets and primary measure

For workload *w*, seed *s*, and progress fraction *f* ∈ {0.05, 0.10, 0.15, 0.20, 0.25}, let *m*_*w,s*,0_ be the common initial validation metric and let 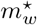 be its mathematical ideal: one for accuracy and perplexity, and zero for error metrics. The target is

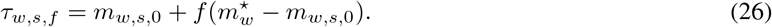

We record the first update at which the target is reached. An unreached target is assigned the full update budget *B*_*w*_ for the restricted-time calculation, while its non-reach is also recorded separately.

For method *a*, define the target-averaged, budget-normalized restricted time

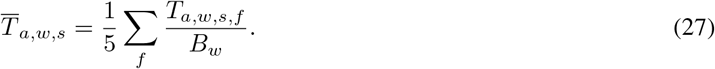

For comparator setting *c*, the primary measure is

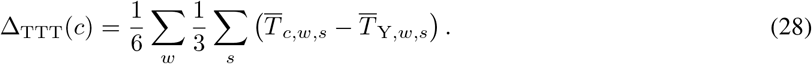

Y denotes YamAdam in Eq. (28). Δ_TTT_ is the primary result of this study. A positive value means that YamAdam reached the predefined targets in fewer optimizer updates on the equal-workload average.

The W/T/L columns summarize the 18 workload–seed cases in each row after the five targets have been averaged: W means YamAdam was faster, T means the two restricted times were equal, and L means the comparator was faster. The B/Y/X/N columns summarize the 90 individual target observations in each row: B means both reached the target, Y means only YamAdam reached it, X means only the comparator reached it, and N means neither reached it.

## 5 Results

Table 2 reports all 30 fixed settings. The bold numeric column is the primary measure Δ_TTT_; positive values favor YamAdam.

All 30 settings had a positive Δ_TTT_, ranging from +0.003526 to +0.210524. Across the 540 setting–workload–seed cases, YamAdam was faster in 504, tied in 14, and slower in 22. Eighteen settings favored YamAdam in all 18 workload–seed cases; the other 12 contained at least one comparator-faster case. Across 2,700 target observations, both methods reached 2,634 targets and only YamAdam reached 66. There were no comparator-only or neither-reached observations.

## 6 Discussion

### 6.1 Target attainment without user tuning

Every reported comparator row represents one fixed setting that a user could choose. YamAdam is used without optimizer settings supplied by the user and had a positive Δ_TTT_ against all 30 rows. This result is stronger than a comparison with one fixed setting because it includes three distinct fixed settings for every comparator family and does not select a favorable family winner after observing the outcomes.

The observed behavior is consistent with the roles of the three main parts of the method. Coordinate-wise moments provide local scale adaptation; historical-only projection prevents stale motion from opposing the current gradient without discarding the current contribution; and one-time calibration plus dimensionless directional statistics determines a global distance without a user-specified learning rate. This interpretation explains the design but does not isolate the causal contribution of each component.

### 6.2 Average and individual comparisons

Δ_TTT_ gives every workload equal weight, whereas W/T/L shows how consistently the result holds across individual workload–seed cases. The 22 losses show that YamAdam was not uniformly fastest. Conversely, the absence of comparator-only target reaches indicates that the positive mean differences were not obtained by sacrificing reach within the evaluated budgets. Both views are needed: the primary average answers the cross-workload question, and the case counts show its limits.

## 7 Conclusion

We introduced YamAdam, an adaptive optimizer that requires no user-specified optimizer hyperparameters. It combines short-memory Adam moments with both decay factors fixed at 1*/*2, historical-only projection, one-time scale calibration, and later scaling from dimensionless directional information. Across six heterogeneous workloads and three seeds, every one of 30 fixed comparator settings had a positive equal-workload Δ_TTT_, and YamAdam was faster in 504 of 540 setting–workload–seed comparisons. These findings support YamAdam as a promising optimizer when workload-specific optimizer tuning is undesirable. The YamAdam implementation is available at https://github.com/caramelazuki/YamAdam.

